# Heat stress induces a developmental shift from type-V to type-IV trichome dependent on jasmonate signaling in tomato

**DOI:** 10.1101/2023.07.28.551053

**Authors:** Robert Säbel, Alejandro Brand, Nick Bergau, Gerd U. Balcke, Frank Syrowatka, Mandy Püffeld-Raorane, Bettina Hause, Alain Tissier

## Abstract

Cultivated tomato (*Solanum lycopersicum*) and related wild species develop several types of trichomes, both glandular and non-glandular, on their aerial parts. Among these, type-IV trichomes are responsible for the synthesis and secretion of acylsugars, which act as defense compounds against herbivores. In contrast to related wild species such as *S. pennellii*, type-IV trichomes are present only in the juvenile stages of cultivated tomato plants and absent in later stages of development. By submitting tomato plants to high temperatures during the day (37 °C), we observe that non-glandular type-V trichomes are replaced by type-IV trichomes. This is accompanied by a massive increase in acylsugar production. On the other hand, heat treatment does not affect type VI-trichomes, which produce mono- and sesquiterpenes, but the production of monoterpenes is increased while that of sesquiterpenes is suppressed. Furthermore, tomato *jai1* mutants deficient in jasmonate (JA) perception do not exhibit this developmental switch from type-V to type-IV trichomes. The implication of JA signaling in this process was further supported by an increase in JA-isoleucine and in the expression of genes involved in JA-signalling within hours of heat stress application. These results establish a unique system to study how environmental factors affect developmental fate decisions in plants while opening opportunities to understand mechanisms controlling type-IV trichome initiation and development.

## Introduction

Glandular trichomes (GTs) are epidermal outgrowths that act as factories of plant specialized metabolites. These substances play an essential role in the interaction of plants with their environment, for example as defense against herbivores, attraction of pollinators or protection from abiotic stresses (Wagner et al., 2004). In the cultivated tomato, *Solanum lycopersicum*, eight different types of trichomes have been described, namely the non-glandular type-II, III, V and VIII, and the glandular type-I, IV, VI and VII (Luckwill, 1943; Glas et al., 2012). Between tomato GT types, there is variation in the cell architecture and the type of molecules they produce. The gland cells of type-VI GTs are known to produce and store volatile organic compounds (VOCs) including mono- and sesquiterpenes (Schilmiller et al., 2010; Bleeker et al., 2012; Tissier et al., 2017; Zhou and Pichersky, 2020; Zabel et al., 2021). On the other hand, the capitate type-I and IV GTs synthesize and secrete sticky substances known as acylsugars (AS) (Li et al., 2014; Nakashima et al., 2016). AS consist of sugar cores, usually sucrose or glucose, esterified with fatty acyl chains at different positions (Schilmiller et al., 2012; Ning et al., 2015). The structural variations of these sugar esters across *Solanaceae* results in an extensive diversity of specialized metabolites with unique chemical properties and biological functions (Moghe et al., 2017; Fan et al., 2019; Fan et al., 2020). In aerial parts of the plants, AS function as insect traps due to their viscosity (Feng et al., 2022), as a protectant against fungal pathogens (Luu et al., 2017), or as a meal for herbivores that after being metabolized, makes them more attractive for predators (Weinhold and Baldwin, 2011). In the roots, AS are exuded in the systemic side of a split-root system when the local side is interacting with specific microbial colonizers, although further research is needed to elucidate their specific function (Korenblum et al., 2020). Thus, AS confer multiple benefits to plants, encompassing not only protection but also communication with the surrounding environment.

In the aerial parts, AS are mainly produced in type-IV GTs, which consist of one basal cell, several stalk cells and one secretory cell at the end of the stalk (Tissier, 2012). Wild tomato species such as *S. pennellii* and *S. galapagense* exhibit a high density of type-IV GTs and AS levels in these species can represent up to 20% of the leaf dry biomass (Fobes et al., 1985; Schilmiller et al., 2012). The high concentration and the chemical diversity of AS in wild tomatoes contribute to a broad spectrum of resistance to different types of arthropods including aphids, spider mites, whiteflies, moths and thrips (Rodriguez et al., 1993; Maluf et al., 2010; Mirnezhad et al., 2010; Vosman et al., 2019). In contrast, domesticated tomato has a low AS content correlated with a low density of type-IV trichomes, rendering it more vulnerable to pests and diseases. In *S. lycopersicum*, type-IV GTs are restricted to early stages of the plant development, from cotyledons where they are predominant, followed by the first set of true leaves where their density starts decreasing, until later stages where they are rarely observed or completely absent (Vendemiatti et al., 2017). Therefore, it was proposed that the presence of type-IV trichomes is a juvenile trait in cultivated tomato (Vendemiatti et al., 2017). Additionally, tomato mutants with heterochronic alterations display type-IV trichomes in fully-grown plants (Vendemiatti et al., 2017). One interesting observation is that the decline in type-IV trichomes inversely correlates with the progressive increase of type-V trichomes, which have the same structure as type-IV but lack the glandular cell at the tip. Hence, it is tempting to speculate that the absence of this type of GTs in adult tomato leaves corresponds to a gradual shift in trichome fate during plant development.

Several efforts were conducted to identify regulators of the type-IV GTs/AS trait using QTL mapping of recombinant inbred lines obtained from crosses between wild and cultivated tomato species (Resende et al., 2002; Firdaus et al., 2013; Vosman et al., 2019). However, the characterization of the causal gene(s) in the identified chromosomal regions remains elusive. Recently, it was shown that the introgression of three alleles from *S. galapagense* into the model tomato Micro-Tom could increase type-IV trichome density in late stages of development, but it was insufficient to subsequently augment the AS quantities (Vendemiatti et al., 2022). These findings provide evidence that type-IV trichome initiation and AS biosynthesis are uncoupled processes that could be governed by distinct regulatory mechanisms. Nevertheless, identifying the genes controlling this juvenile trait represents the first step in the engineering of insect-resistant tomatoes.

The herbivore and wound-induced phytohormone jasmonic acid (JA) and its conjugates play an important role in trichome initiation. JA is synthesized from fatty acids released from the plastid membranes that after several enzymatic reactions are converted into (+)-7-*iso*-jasmonoyl isoleucine (JA-Ile), the bioactive from of JA (Schaller and Stintzi, 2009; Wasternack and Hause, 2013). Upon elicitation, JA-Ile is perceived by the F-box protein CORONATINE INSENSITIVE1 (COI1), a component of the SCF^COI1^ complex that targets the JASMONATE ZIM-domain (JAZ) proteins for proteasome degradation, the latter being repressors of JA signaling (Chini et al., 2007; Thines et al., 2007; Fonseca et al., 2009). Once JAZ are degraded, transcription factors such as MYC2 (and other members of the basic helix-loop-helix family) are released and mediate the activation of JA responsive-genes (Fernández-Calvo et al., 2011; Wasternack and Hause, 2013). In tomato, the *jasmonic acid-insensitive 1* (*jai1*) mutant, which carries a deletion within the ortholog of Arabidopsis *COI1*, displays defects in pollen and seed development and a significant reduction in type-VI trichome density, highlighting the function of JA/COI1 signaling in trichome initiation and development (Li et al., 2004). COI1 can interact with the JAZ2 repressor, which is a negative regulator of trichome development and terpene biosynthesis (Thines et al., 2007; Yu et al., 2018; Hua et al., 2021). Moreover, the application of JA or methyl jasmonate (MeJA) induces glandular and non-glandular trichome formation as well as increases the length of trichomes (Boughton et al., 2005; Escobar-Bravo et al., 2017; Chen et al., 2018; Hua et al., 2021). Although the connection between JA signaling and type-VI trichomes has been extensively reported, little is known about the regulation of type-IV trichome development in tomato and the possible link to JA or other stress responses.

In the present study, we report that heat stress (HS) application induces a shift from type-V to type-IV trichome development in later stages of tomato growth. We show that besides promoting type-IV trichome initiation, elevated temperature boosts acylsugar biosynthesis exponentially. Although no change in type VI-trichome density was observed, HS conditions modified the volatiles composition and content. Finally, we reveal that the *jai1* mutant does not exhibit the type-IV trichome phenotype in HS, supporting the role of jasmonate signaling in the determination of the trichome fate of the type-IV/V lineage.

## Results

### Heat stress induces type-IV trichomes in cultivated tomato

We exposed plants of *S. lycopersicum* cv. Moneymaker to heat (37 °C; HS) during the day for 3 weeks, after 1 week of germination under control conditions (25 °C; CN; **Fig.1A**). Counting type-IV and type-V trichomes on the edge of the third and fourth leaf from heat-treated plants (**Fig. 1B**) revealed a strong increase in type-IV trichomes compared to plants that were grown under moderate temperatures (**Fig.1C**). Simultaneously, the number of type-V trichomes decreased drastically, resulting in an inverted ratio of type-IV to type-V trichomes in heat-treated plants. The sum of type-IV and V trichomes was not significantly altered, only the trichome type was affected by heat.

**Figure 1.**
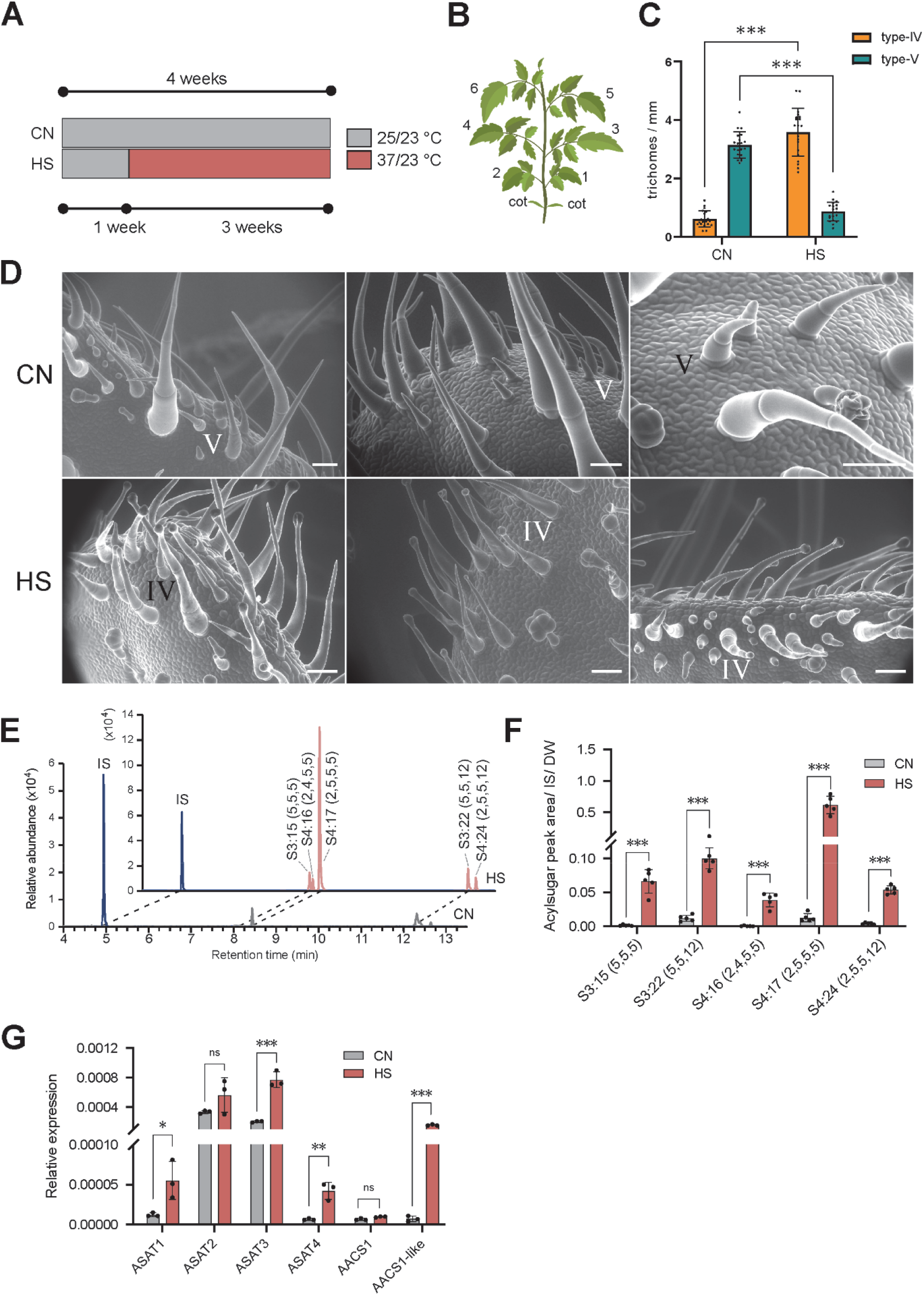
Effect of heat stress on type-IV and -V trichomes and acylsugar (AS) content. (A) Scheme of growing conditions for control (CN) and heat stress plants (HS). (B) Schematic drawing of a tomato plant with leaf numbers (C). Number of type-IV and type-V trichomes on the edge of leaflets from the third and fourth leaf (*n* = 6 biological replicates) in CN and HS conditions. (D) Scanning electron microscopy of leaflets from the third leaf. Scale bar = 50 µm. (E) Representative LC-MS chromatogram displaying AS peaks in CN and HS. (F) Quantification of representative AS using ultra-performance liquid chromatography with mass spectrometry (UPLC-MS). Chromatogram peak areas were normalized to an internal standard (IS) and leaf dry weight (DW). (G) Relative expression levels of selected genes involved in AS biosynthesis (*n* = 3 biological replicates). ASAT, acylsugar acyltransferase; AACS, acylsugar acyl-CoA synthetase. Error bars indicate standard deviation. \**P* < 0.05, \*\**P* < 0.01, \*\*\**P* < 0.001; using Student’s *t*-test.

Images produced with an environmental scanning electron microscope (ESEM) on young, developing leaflets that underwent the heat treatment, illustrate the shift in trichome type (**Fig. 1D**). In addition, these images show that the single trichome secretory cell at the tip of the trichome, indicative of a type-IV trichome, is established at very early stages of trichome development, leading to the conclusion that the trichome type fate is determined already upon the initiation of a trichome. Young, emerging type-V trichomes lack such a head cell and exhibit already their typical pointy tip. Thus, the inversion in the ratio of type-IV/type-V is not due to a developmental switch after trichome initiation, but rather to a switch upon trichome initiation to engage into type-IV instead of type-V development.

To further investigate this phenomenon, acylsugars (AS), the main products of type-IV trichome, were analysed by ultra-performance liquid chromatography-mass spectrometry (UPLC-MS) comparing surface extracts of plants from control and heat stress conditions (**Fig. 1E**). AS consist of a sucrose or glucose moiety esterified by acyl chains of short to medium length at different positions (Kroumova et al., 2016). In tomato, the sugar is sucrose and the acyl chains are mostly C2, branched C4 and C5 (*iso* and *anteiso*), and C12 (Ghosh et al., 2014; Kroumova et al., 2016). Consistent with the elevated type-IV trichome numbers, the AS content was increased in plants under heat stress (**Fig. 1F**). Remarkably though, this increase was much stronger than the increase in type-IV trichomes. While the type-IV trichomes were five to six times more abundant after HS, the AS content increased between eight to 76 times depending on the individual AS. This hints at an additional process that leads to higher AS productivity. The AS composition was not affected qualitatively by heat stress.

The last steps in the biosynthesis of AS in tomato, the esterification of acyl chains to the sugar backbone, are catalysed by acylsugar acyltransferases (ASATs) (Kim et al., 2012; Schilmiller et al., 2015; Fan et al., 2016). Acylsugar acyl-CoA synthetases (e.g. AACS1) prime the acyl chains for esterification by activating the carboxylic acid group via conjugation to CoA (Fan et al., 2020). All the respective genes are preferentially expressed in the glandular cell of type-IV trichomes (Fan et al., 2020). To gain insight into the enhanced AS productivity in heat stressed tomato plants, we measured the transcript levels of different ASAT and AACS encoding genes by quantitative RT-PCR (**Fig. 1G**). All the analysed genes except *ASAT2* and *AACS1* were significantly upregulated in HS. Nevertheless, those two genes showed an upward trend too. The expression of most genes increased between three to six fold, with the exception of *AACS1*-*like*, whose expression was more than 22 times stronger under heat.

### Longer heat stress leads to stronger phenotype

We next sought to determine when the HS should start and how long it should last to induce the type-IV trichome phenotype on the third leaf using AS measurements as a proxy. We found that starting at day 7 and applying heat for 4 to 9 nine days did not trigger the type-IV trichome phenotype (**Supp Fig. 1**). We also found that applying HS starting on day 16 was still sufficient to trigger the expression of type-IV trichomes on the third leaf (**Supp Fig. 1**). Next, from day 16 onwards the plants were treated with heat for 2, 3, 4 and 6 days before undergoing a recovery phase at 25 °C until the observation at day 28 (**Fig. 2A-B**). Already after a 2-day heat period there was a significant increase in type-IV trichomes compared to the control plants (**Fig. 2C**). The longer the heat period, the more type-IV trichomes were present, accompanied with a simultaneous decrease of type-V trichome abundance. A heat period of six days was sufficient to trigger more type-IV than type-V trichomes. AS measurements supported the trichome counting (**Fig. 2D**). The longer the heat periods, the higher the AS content in the leaf extracts. Two days of HS doubled the overall AS abundance compared to CN. Four days of heat increased the AS nearly ten times. This increase again cannot be attributed only to the type-IV trichome induction by heat. We also submitted several tomato cultivars to heat treatment and observed a similar trichome switch phenotype and increase in AS production, albeit to variable extent in some cultivars (e.g. LA1420 and Criollo) (**Supp. Fig. 2 and 3**)

**Figure 2.**
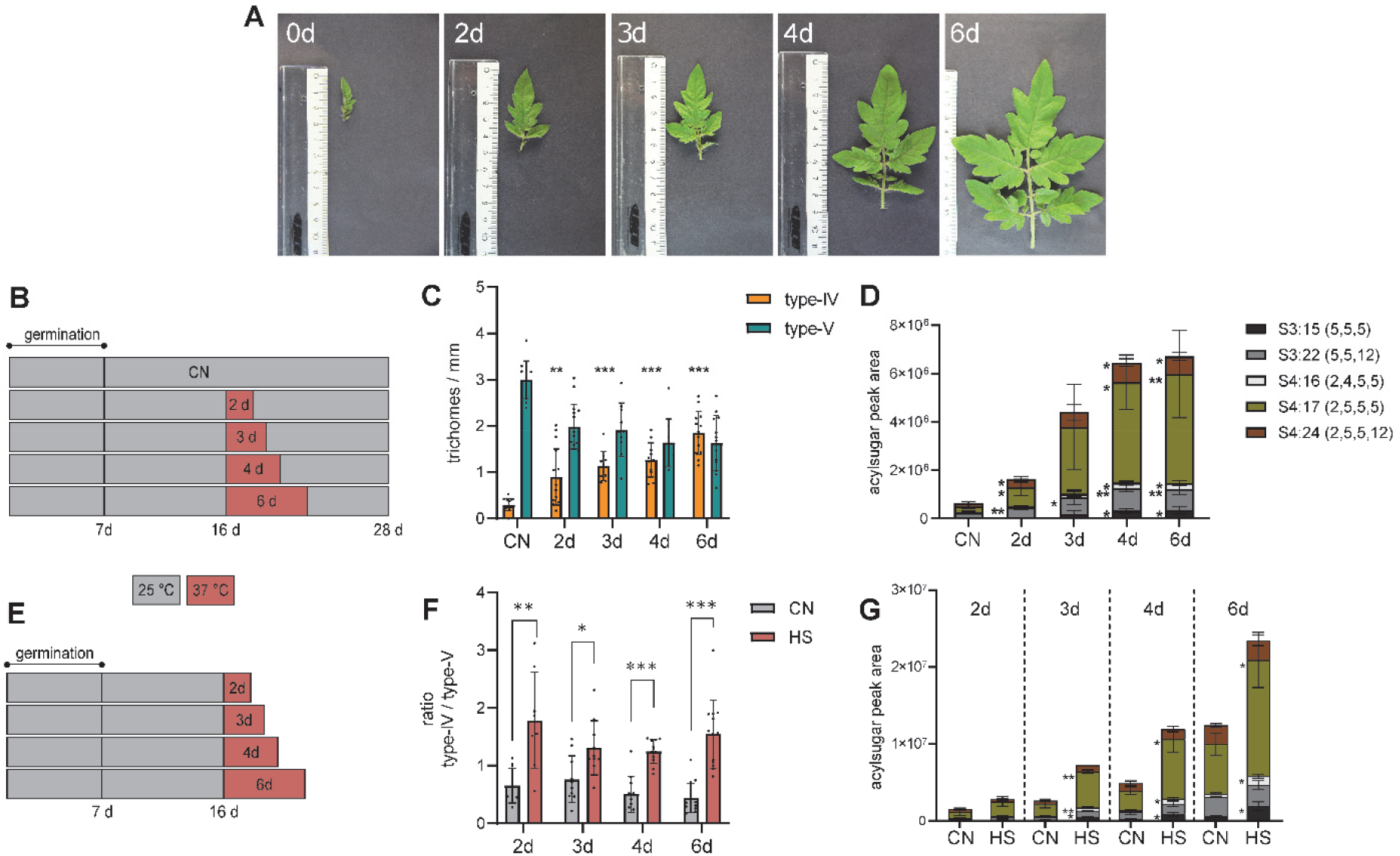
Influence of length and timing of heat stress. (A) Photos of third leaves from Moneymaker plants after 16 d post sowing (0 d, when heat treatment started) to 22 d post sowing. (B) + (E) Schemes of growing conditions with different lengths of heat treatment followed (B) or not (E) by a recovery phase. (C) Number of type-IV and type-V trichomes on the edge of leaflets from the third leaf after different heat periods as shown in (A) (n = 3 biological replicates). (F) Ratio of type-IV and type-V trichomes on the edge of leaflets from the third leaf after different heat periods as shown in (E). (D) and (G) Quantification of AS from plants grown as in (B) and (E) respectively. Error bars indicate standard deviation. \**P* < 0.05, \*\**P* < 0.01, \*\*\**P* < 0.001; using Student’s *t*-test. *P* values indicated in relation to corresponding CN.

**Figure 3.**
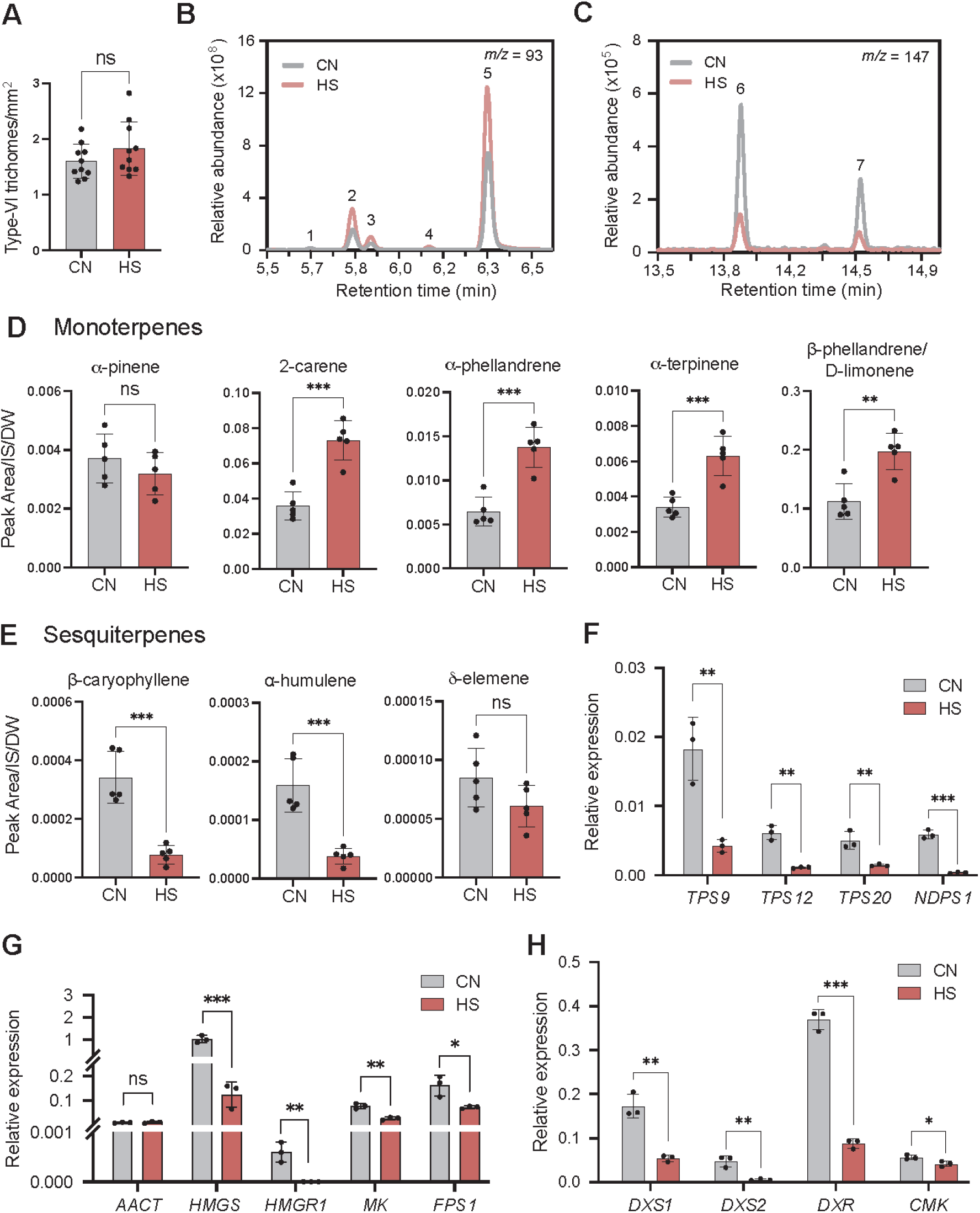
Effect of heat stress on type-VI trichomes and terpenoid profiles. (A) Type-VI trichome number in tomato plants after three weeks of growth in CN and HS conditions (*n* = 10 biological replicates). Representative chromatograms displaying monoterpene peaks (B) and sesquiterpene peaks (C) in CN and HS. 1: α-pinene, 2: 2-carene, 3: α-phellandrene, 4: α-terpinene, 5: β-phellandrene+limonene, 6: β-caryophyllene, 7: α-humulene. Quantification of monoterpenes (D) and sesquiterpenes (E) using gas chromatography-mass spectrometry (GC-MS) (*n* = 5 biological replicates). Chromatogram peak areas were normalized to internal standard (IS) and to leaf dry weight (DW). Relative expression levels of terpene synthases (F) and selected genes involved in the mevalonate (G) and the methylerythritol phosphate (H) pathways (*n* = 3 biological replicates). *TPS*, terpene synthase; *NDPS1*, *Neryl-diphosphate synthase 1*; *AACT*, *Acetyl-CoA C-acetyltransferase*; *HMGS*, *hydroxymethylglutaryl-CoA synthase*; *HMGR1*, *3-hydroxy-3-methylglutaryl-coenzyme A reductase 1*; *MK*, *mevalonate kinase*; *FPS1*, *farnesyl diphosphate synthase*; *DXS*, *deoxy-xylulose phosphate synthase*; *DXR*, *deoxy-xylulose phosphate reductoisomerase*; *CMK*, *4-(cytidine 5’-diphospho)-2-C-methyl-D-erythritol kinase*. Error bars indicate standard deviation. \**P* < 0.05, \*\**P* < 0.01, \*\*\**P* < 0.001; using Student’s *t*-test.

In an additional HS time series, trichome counting and AS measurements were performed directly following the different heat periods instead of at day 28 (**Fig. 2E**). With this setup, the immediate impact of heat stress on the trichomes can be observed. The leaflets are in early developmental stages and expand drastically over the span of the heat periods (**Fig. 2A**), resulting in a decreasing trichome density over time both on the edge and the inner surface of the leaflet. Considering this, the trichome count is shown as a ratio between type-IV and type-V trichomes rather than in total numbers. Additionally, corresponding controls to each time point were included for the trichome count and the AS measurements. In the controls, the ratio of type-IV to type-V trichomes remained under 1, independently of the time point (**Fig. 2F**). After two days, the ratio of type-IV to type-V increased from 0.5 in control conditions to almost 2 under heat. A similar ratio was observed for all the time points up to six days of heat. The AS content in the controls increased in an exponential fashion over the 6 days (**Fig. 2G**). This indicates that in the early stages of leaf development, AS production is not immediately active upon trichome formation. In heat, the AS levels increased even further, by 1.8 to 2.7 times compared to the corresponding controls. This is in the same range as the increased abundance of type-IV trichomes, an observation differing from the previous experiments, where the measurements were made at day 28 and where the fold-increase in AS production exceeded that of type-IV trichome.

### Effect of heat stress on type-VI trichome development and productivity

Type-VI trichomes are the most abundant type of glandular trichomes present in cultivated tomato *S. lycopersicum*. They produce large amounts of volatile organic compounds (VOCs) that are stored in a dedicated cavity surrounded by four glandular head cells (Bergau et al., 2015). We examined the impact of HS on type-VI trichome development and the VOCs productivity using the same experimental set-up as before (**Fig. 1A**). First, trichomes were counted over the surface of leaf four and five (**Fig. 1B**) of plants growing in HS for three weeks. No significant differences were found in trichome density compared to the respective control (**Fig. 3A**). However, contrasting changes in the VOC profiles were observed (**Fig. 3B-C**). While most of the monoterpenes increased under heat, the sesquiterpenes showed a significant reduction (**Fig. 3D-E**). In *S. lycopersicum*, sesquiterpenes (C15) originate from the cytosolic mevalonate (MEV) pathway and monoterpenes (C10) from the plastidic methylerythritol phosphate (MEP) pathway. We hypothesized that an increase or decrease in the accumulation of VOCs may be supported by changes in the transcriptional activity of genes requited for their biosynthesis. RNA from leaflets was extracted and the relative expression of relevant genes was measured by qRT-PCR. Expression levels of *Terpene Synthase 9* (*TPS9*) and *TPS12*, whose main products are germacrene and β-caryophyllene/α-humulene respectively (Zhou and Pichersky, 2020), were significantly reduced in heat (**Fig. 3F**). Concomitantly, genes encoding enzymes of the MEV pathway such as *hydroxymethylglutaryl-CoA synthase* (*HMGS*), *3-hydroxy-3-methylglutaryl-coenzyme A reductase 1* (*HMGR1*), *mevalonate kinase* (*MK*) and *farnesyl diphosphate synthase* (*FPS1*) also showed lower expression levels at elevated temperature (**Fig. 3G**). Surprisingly, the levels of *TPS20*, responsible for synthesizing the major monoterpenes in type-VI GTs including 2-carene, α-terpinene, α-phellandrene and β-phellandrene (Zhou and Pichersky, 2020), as well as of *Neryl-diphosphate synthase 1* (*NDPS1*), that catalyzes their immediate precursor (Schilmiller et al., 2009), were also downregulated (**Fig. 3F**) . At the same time, genes that determine the flux through the MEP pathway, such as *deoxy-xylulose phosphate synthase* (*DXS1*), *DXS2*, *deoxy-xylulose phosphate reductoisomerase* (*DXR*), *4-(cytidine 5’-diphospho)-2-C-methyl-D-erythritol kinase* (*CMK*) exhibited a significant reduction in transcript levels after a long-term heat exposure (**Fig. 3H**).

Unlike tomato, *Arabidopsis thaliana* has only one type of trichomes, which consist of a single cell that is branched. To check whether the developmental switch we observed in tomato is specific to glandular trichomes, we submitted Arabidopsis plants to different temperature regimes, thereby applying heat stress over several days. Counting trichome numbers on rosette leaves 8 and 13, we did not find significant differences in the density or morphology of trichomes (**Suppl. Fig. 4)**.

### Jasmonate signalling and trichome development under heat stress

In tomato, several studies demonstrate the key role of jasmonic acid (JA) (Boughton et al., 2005; Escobar-Bravo et al., 2017; Chen et al., 2018) and JA signaling in trichome development (Chalvin et al., 2020). We thus sought to explore whether the induction of type-IV trichomes in heat stress could be mediated by JA. The *jasmonic acid-insensitive 1* (*jai1*) mutant is defective in JA signaling and displays reduction in trichome density in several organs of the plant (Li et al., 2004). Since *jai1* mutants are female sterile, homozygous plants were obtained from heterozygous (*+/jai1*) plants. Seeds of tomato *S. lycopersicum* cv. Castlemart harboring the *jai1* mutation were sown on soil under control temperature conditions (CN, 25 °C 16h day/23 °C 8h night) and after ten days the seedlings were genotyped by PCR. Ten plants per genotype (+/+, +/*jai1*, and *jai1*/*jai1*) were either maintained in CN or transferred to heat stress (HS, 37 °C 16h day/23 °C 8h night) for another 20 days. As in previous experiments, wild-type (WT) and *+/jai1* plants showed a significant increase in type-IV trichomes, and a decrease in type-V trichomes (**Fig. 4A**) under HS compared to CN. However, in homozygous *jai1/jai1* plants, there was no significant increase in type-IV trichomes (**Fig. 4A**). Aligned with the lack of type-IV trichomes, no significant increase in AS content was detected in the *jai1* mutant (**Fig. 4B**). Regarding type-VI trichomes, high variation in trichome numbers was observed among the different genotypes and a slight reduction was observed in the *jai1* mutants at elevated temperature (**Fig. 4C**). Moreover, an increase in the major monoterpenes was observed in all genotypes albeit to a reduced extent in +/*jai1* and *jai1/jai1* genotypes, while the sesquiterpenes were reduced regardless of the genotype (**Fig. 4D-E**).

**Figure 4.**
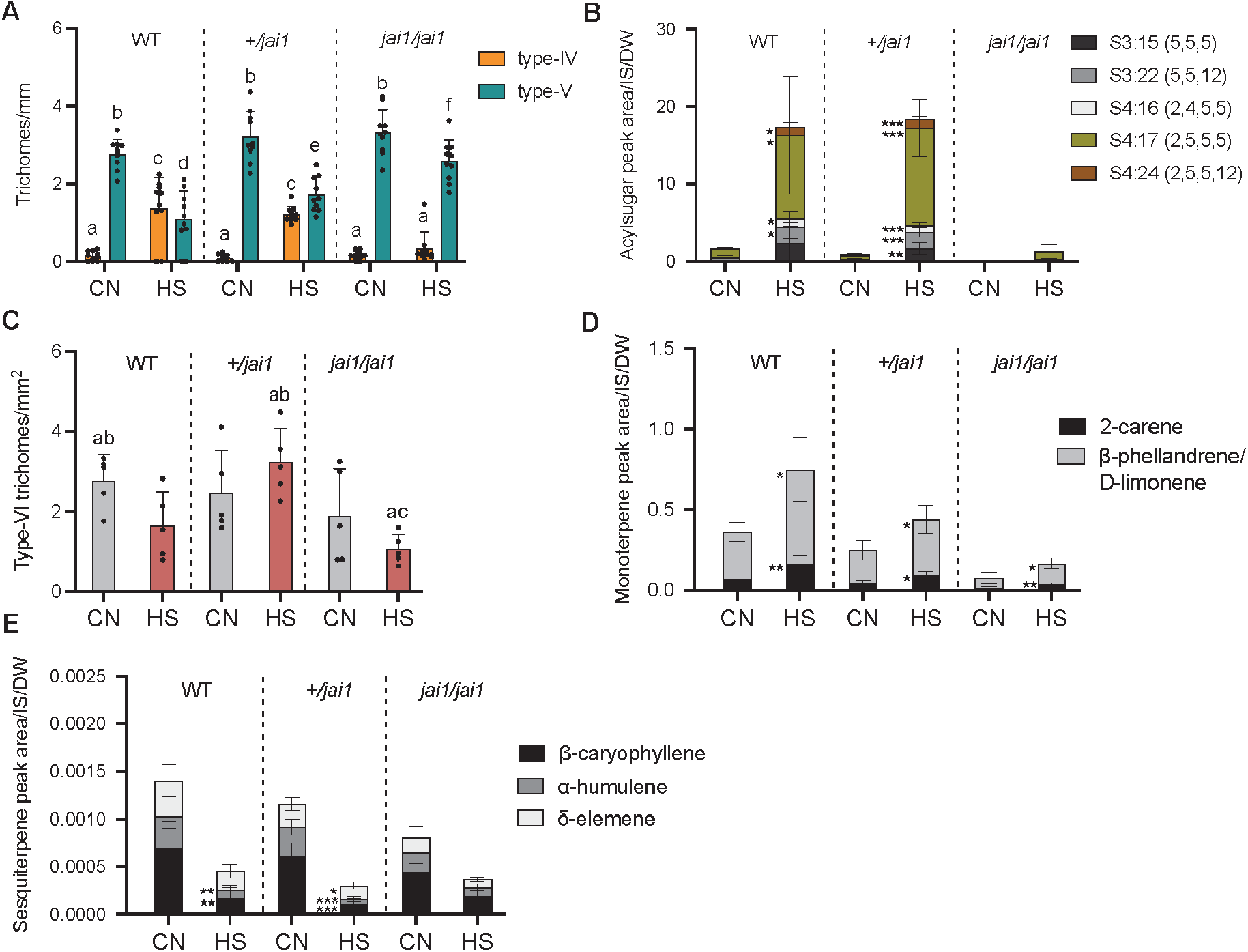
Heat stress does not induce type-IV trichome initiation in mature leaves of the *jasmonic acid-insensitive1* (*jai1*) mutant. (A) Type-IV/V trichome counting on leaves of wild type (WT), heterozygous (*+/jai1*) and homozygous (*jai1/jai1*) plants after three weeks of growth under CN and HS conditions (*n* = 10 biological replicates). (B) Acylsugar quantification by UPLC-MS. (C) Type-VI trichome number (*n* = 5 biological replicates). The only significant differences are between *jai1/jai1* HS (ac) and WT CN (ab) and between *jai1/jai1* HS (ac) and *+/jai1* HS (ab). (D-E) Estimation of the most abundant monoterpenes (D) and sesquiterpenes (E) produced by type-VI trichomes using GC-MS. Chromatogram peak areas were normalized to internal standard (IS) and to leaf dry weight (DW) (*n* = 5 biological replicates). Error bars shows standard deviation. Letters indicate two-way ANOVA (*P* < 0.05, using Tukey’s test). Pairwise comparisons were done using Student’s *t*-test \**P* < 0.05, \*\**P* < 0.01, \*\*\**P* < 0.001.

### Jasmonate hormone levels and expression of JA signaling genes under heat stress

To gain further insights into the JA-signaling dependency of the heat-induced tomato trichome phenotype, we measured the levels of OPDA, JA and jasmonoyl-isoleucine (JA-Ile) in control and heat stressed plants from WT, +/*jai1* and *jai1*/*jai1* after three weeks of HS exposition (**Fig. 5A**). OPDA levels were lower in +/*jai1* and *jai1*/*jai1* compared to WT, but there was no difference between HS and CN conditions. JA levels showed no difference, regardless of genotype or condition. There was a trend for higher JA-Ile levels in HS plants, but this was not significant. Thus, no striking differences in JA-related metabolites were observed after long-term exposure to heat. However, our heat stress time series showed that a two day-exposure to heat is sufficient to trigger the trichome development switch (**Fig. 2B**), indicating that the required JA-signaling events are likely to occur shortly after exposure to heat. We thus chose to measure JA-related metabolites under HS and CN conditions within the first six hours of heat (**Fig. 5B**). We observed that OPDA levels increased significantly, reaching a peak after 2 h, while JA and JA-Ile exhibited a gradual increase starting at 30 min after heat application (**Fig. 5C**). We then measured the expression of genes involved in JA-signaling by qRT-PCR within the same 6 h time frame (**Fig. 5D**). *Heat shock protein 70* (*HSP70*) involved in heat stress response was used as a marker to monitor the heat treatment and showed a rapid increase in transcript levels that peaked at 30 min. A similar pattern was observed for *JAZ2*, albeit with much lower expression levels and *COI1* showed a moderate increase after 15 and 30 min of HS as well. *JAZ4* showed a gradual increase with a maximum at 6 h and *MYC2* displayed significantly increased expression at 6 h. Thus, changes in JA and related metabolites, as well as alterations in gene expression of JA-signaling pathway genes were detected after a short time exposure to elevated temperature.

**Figure 5.**
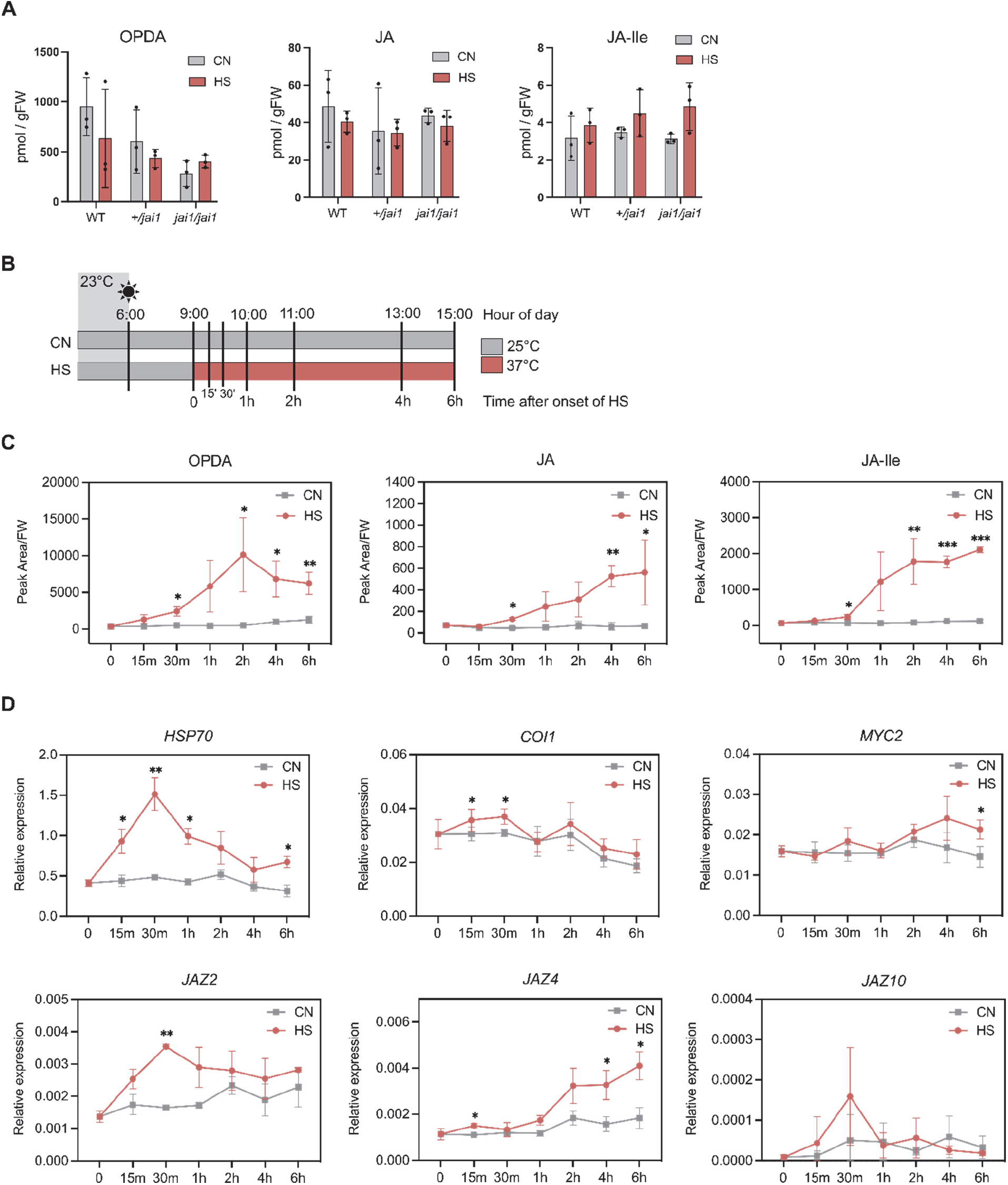
Long- and short-term JA response to heat stress. (A) Phytohormone levels in leaves of plants after three weeks of growth under CN and HS conditions (*n* = 3 biological replicates). (B) Schematic representation of the short HS time series indicating the time points when leaf tissue was collected for phytohormone and gene expression measurements. (C) Phytohormone levels after short HS exposition. Chromatogram peak areas were normalized to leaf fresh weight (FW) (*n* = 3 biological replicates). OPDA, *cis-* 12-oxo-phytodienoic acid; JA, jasmonic acid; JA-Ile, (+)-7-*iso*-jasmonoyl isoleucine. (D) Relative expression of JA signaling genes after HS exposure. *Heat shock protein 70* (*HSP70*) was used as a control of the heat treatment. Error bars display standard deviation. Pairwise comparisons were done using Student’s *t*-test \**P* < 0.05, \*\**P* < 0.01, \*\*\**P* < 0.001.

## Discussion

### Heat stress induces type-IV trichome initiation in tomato

We uncovered here a developmental phenotype induced by heat stress, which to the best of our knowledge had gone unnoticed before. It is worth noting that in related wild tomato species, especially *S. pennellii*, type-IV trichomes are present at high density in mature leaves constitutively and produce large amounts of AS. The fact that cultivated tomato can develop type-IV trichomes for example on cotyledons or in young leaves (Vendemiatti et al., 2017), suggests that upon domestication the ability to develop high numbers of type-IV trichomes in mature leaves has become repressed.

Our observations that heat stress induces a shift in developmental fate of epidermal cells in tomato provide an illustration of how environmental conditions can affect developmental decisions. Such developmental fate decisions dependent on the environment are present in fungi. The yeast *Saccharomyces cerevisiae* for example, which is normally a unicellular organism, switches to a filamentous growth when starved for nitrogen (Gimeno et al., 1992), a decision which is dependent on a MAP kinase signaling cascade (Liu et al., 1993; Roberts and Fink, 1994). Such alterations in developmental fate decisions are also known in plants. For example, a number of aquatic plants like *Callitriche heterophylla* develop two strikingly different leaf morphologies, for the submerged and emerged leaves respectively (Deschamp and Cooke, 1983). Interestingly, these can be manipulated by various treatments, including temperature, gibberellic acid or a change in osmolarity (Deschamp and Cooke, 1983). Unlike *C. heterophylla*, tomato is a genetically and molecularly tractable model species, making it possible to elucidate the mechanisms underlying this developmental fate decision process.

Unlike Arabidopsis, which has only one major type of non-glandular trichome, tomato and many other plant species such as *Artemisia annua*, have multiple trichome types including glandular and non-glandular types. There is currently very little information and knowledge on what determines the fate epidermal cells to generate one specific type of trichome. Our observations raise the question whether this developmental switch is specific to glandular trichomes. Our results with Arabidopsis, did not reveal significant difference in the density or morphology of the trichomes (**Supp. Fig. 4**). This would suggest that the morphological switch we observe in tomato is specific to glandular trichomes, although a much broader survey would be required to support this statement.

### Heat stress induces type-IV trichomes in a COI1 dependent manner

Jasmonates are well known as stress phytohormones involved in the defense response to herbivores and in the regulation of several growth and developmental aspects (Ahmad et al., 2016; Howe et al., 2018) but little is known about the connection between JA signaling and the plant adaptive responses to elevated temperatures. Different studies have shown in Arabidopsis that short regimes of heat stress increase the levels of JA and induce the expression of JA biosynthesis and signaling-related genes such us *OPR3*, *MYC2* and JAZ proteins (Clarke et al., 2009; Cortijo et al., 2017; Tian et al., 2020; Agrawal et al., 2022). JA-mediated signaling pathways appear to play a role in response to warm temperatures in other plant species as well. For example in wheat (*Triticum aestivum*) the silencing of *OPR3* leads to a reduction in thermo-tolerance (Tian et al., 2020). In tomato, it was reported that elevated temperature enhances the JA-dependent wound response due to the accumulation of the COI1 receptor (Havko et al., 2020). Consistent with previous results reported for Arabidopsis, our data show that HS triggers the biosynthesis of JA and JA-related metabolites within 30 min to 1 hr, leading to the activation of the JA signaling pathway, without wounding of the plant tissue (**Fig. 5**). We did not observe such changes in mature leaf tissues after a long-term exposure to HS, suggesting that after a period of acclimatization to high temperature, levels of JA and JA-related metabolites return to background values. Altogether, our results provide strong evidence that induction of JA and JA-signaling in developing leaves play a critical role in triggering the developmental switch from type-V to type-IV trichomes. The fact that that application of methyl jasmonate (MeJA) cannot induce by itself type-IV trichome formation in the tomato cultivar Moneymaker (Escobar-Bravo et al., 2016) indicates that there are other factors triggered by high temperatures besides JA that regulate trichome fate.

In addition to the trichome type switch, the massive increase in AS biosynthesis stands out from the heat stress treatment and proportionally exceeds the increase in type-IV trichome numbers. This shows that heat has two distinct effects, one on the development of trichomes and the other on the increased productivity of AS. Our data indicate that the latter is also JA-signaling dependent (**Fig. 2B**) and most likely leads to the induction of AS biosynthesis genes as shown in **Fig. 1G**. Since AS are secreted, it is also possible that putative AS transporters are induced by heat. However, although it is likely that these transporters do exist, as suggested by a recent study (Vendemiatti et al., 2022), they are not known and therefore we could not test this hypothesis.

AS deficient mutants of *Nicotiana benthamiana*, besides being more susceptible to insects, have a higher water loss and higher leaf temperatures compared to the wild type (Feng et al., 2022). Therefore, it is possible that under heat stress these sticky compounds act as wax-like protectors to prevent desiccation in addition to their properties as insect immobilizing and toxic agents. In *S. pennellii*, type-IV trichomes are present in high densities over the plant (Glas et al., 2012) and produce large amounts of AS representing up to 20% of the leaf dry weight (Fobes et al., 1985). This species is adapted to arid environments (Dariva et al., 2020), however until now, to our knowledge, no connection was made in this species between AS production and tolerance to drought or heat. Although *S. pennellii* x *S. lycopersicum* introgression lines (Eshed and Zamir, 1995; Ofner et al., 2016) could be helpful to determine the contribution of AS to heat and/or drought resistance, the fact that type-IV trichomes and AS biosynthesis are highly multigenic traits makes it difficult to address this question.

### Changes in volatiles upon heat stress

Heat stress can increase terpenoid emission rates in different plant species containing terpene-storage structures such as glandular trichomes or oil glands (Nagalingam et al., 2023). In tomato, short-term heat stress (HS) enhances the biosynthesis of VOCs including mono and sesquiterpenes, but after the stress period is over, the rate of emission drops down as well as the transcription of the terpene synthases (Pazouki et al., 2016; Nagalingam et al., 2022). In our experimental set up (**Fig. 1A**), wild-type plants grown under long day HS conditions for three weeks exhibited an increase in monoterpene production but a reduction of the sesquiterpenes. Since type-VI trichome density did not change due to high temperatures regimes, the rise in the monoterpene content could be attributed to the increase in metabolic rates. In *S. lycopersicum*, mono- and sesquiterpenes are biosynthesized from the MEP and MEV pathways respectively, that take place in different cell compartments (plastids and cytosol, respectively) (Kortbeek et al., 2016; Balcke et al., 2017). A predominance of monoterpenes over sesquiterpenes in high temperatures could indicate high enzymatic activity and fluxes of precursors towards the biosynthesis of plastid-derived terpenes, which can be retained in the storage cavities of the type-VI trichomes (Tissier et al., 2017). At the same time, the down-regulation of enzymes involved in the MEP and MEV pathways we observe, suggests a negative regulation after acclimatization to heat stress conditions. Both for increases in AS and terpene production, a supply of energy and precursors needs to be secured, in particular sucrose as the main carbon source that must be provided from the leaf to the trichomes (Balcke et al., 2017). A recent report shows that plants of the Mediterranean shrub *Halimium halimifolium* under heat stress reallocate their fixed carbon to respiration and production of VOCs, supporting the role of specialized metabolites in the protection against abiotic stresses (Werner et al., 2020). Thus, the regulation of resource allocation under heat stress conditions in GTs opens new questions that requires further investigation.

In conclusion, the heat-induced phenotype we observed in tomato constitutes a unique opportunity to study the molecular basis of the effect of environment on developmental decisions. Our results also open up opportunities to understand the regulation of type-IV trichome development and in the long term to create tomato varieties with higher type-IV trichomes densities for increased protection not only against insects but possibly also against abiotic stresses.

## Methods

### Plant material and growth conditions

Plants of tomato *Solanum lycopersicum* cultivar Moneymaker, Castlemart and *jai1* mutant in Castlemart background were grown on soil in phytocabinets (CLF Plant Climatics, Emersacker, Germany; model AR-66-L) under long-day (16h light, 8 h darkness) conditions. In control conditions (CN), the daytime temperature was 25 °C and in heat stress conditions (HS) it was 37 °C. In both conditions the temperature at night was set to 23 °C, the relative humidity was 65 % and the light intensity between 150 and 200 pmol _s-1 m2._

For the HS experiments, seeds were germinated and seedlings grown for 7 days in CN conditions. Afterwards seedlings were transferred to HS conditions for three more weeks. The control plants stayed in CN conditions.

For the HS time series experiments (**Fig. 2B+E**), germination and cultivation were performed for 15 days in CN conditions. On Day 16 the heat stress plants were transferred to HS conditions. After different HS periods the plants were transferred back to CN conditions and the phenotyping took place after four weeks (post sowing). Control plants were kept in CN conditions the whole time. In another experiment the phenotyping took place directly after the HS periods without an additional recovery phase.

For the short-term HS time series experiment (**Fig. 5B-C**), seeds of WT Moneymaker were sown on soil in CN conditions. After three weeks of growth, half of the plants were transferred to HS at the indicated time of the day and tissue was collected in the selected time points.

### Trichome counting

For the trichome counting, microscopy pictures of leaflets from the third or fourth leaf were taken with a VHX-6000 microscope in combination with a VH-Z20 T zoom lens (both Keyence, Osaka, Japan) at 30x magnification. Type-IV and type-V trichomes were counted on the edge near the tip of the leaflets. The counting of type-VI trichomes was performed on the adaxial side of 1,5 mm wide strips from leaflets that were cut horizontally to the middle vein and rotated by 90°, or on the whole leaflet. The counting was done manually with the Fiji/ImageJ software. The number of trichomes was normalized to the length of the edge on the picture and the area of the strip or the leaflet respectively.

### Environmental Scanning Electron Microscopy (ESEM)

Scanning electron micrographs of the youngest, still developing leaflet from the fourth true leaf were made with an ESEM XL-30 FEG (FEI/Philips, Eindhoven, The Netherlands) operating in the wet mode. Fresh, unfixed material was used. The gas pressure in the ESEM chamber was 1.5 mBar, regulated by introducing water vapor. Imaging was done with a gaseous secondary electron detector with an acceleration voltage of 12 kV.

### RNA isolation and quantitative RT-PCR

Total RNA was extracted from tomato leaves using the RNeasy© Plant Mini Kit (Qiagen United States, 74904) following the manufacturer’s protocol. For each sample, one microgram of total RNA was treated with DNA-*free*^TM^ DNA Removal Kit (AM1906, Invitrogen) and cDNA was synthesized using the Proto-Script first strand cDNA synthesis kit (New England Biolabs). Quantitative RT-PCR was performed in a CFX Opus Real-Time PCR (Bio-Rad) employing the 5X QPCR Mix EvaGreen (Bio&SELL GmbH) with the following program. Denaturation: 95°C for 15 min; amplification: 40 cycles of 95°C for 15 sec, 58°C for 10 sec and 72°C for 15 sec. At the end of the cycles heating up to 95°C with a heating rate of 0.05°C sec^-1^ was performed to produce melting curves. Target genes were amplified from three biological replicates. The Bio-Rad CFX Maestro© software was used to calculate the Cq values and *S. lycopersicum* Elongation factor 1-alpha (*SlEF1α*) was used in all cases as the reference gene. RT-qPCR primers are listed in Supplemental Table 1.

### Acylsugars and volatiles quantification

Semi-polar metabolites were extracted by placing three freshly punched leaf discs of 1 cm diameter from tomato leaflets in a 2-mL reaction tube containing 1 mL of 80% methanol. Due to the small leaflet size, only one leaf disc was used for the time series experiments. The tube was inverted manually for 2 min. Extracts were transferred to a new 1.5-ml tube, centrifuged for 3 min at 10,000 g, and placed into a glass vial.

Separation of semi-polar metabolites was performed on a Nucleoshell RP18 (2.1 x 150 mm, particle size 2.1 µm, Macherey & Nagel, GmbH, Düren, Germany) using a Waters ACQUITY UPLC System, equipped with an ACQUITY Binary Solvent Manager and ACQUITY Sample Manager (20 µL sample loop, partial loop injection mode, 5 µL injection volume, Waters GmbH Eschborn, Germany). Eluents A and B were aqueous 0.3 mmol/L NH_4_HCOO (adjusted to pH 3.5 with formic acid) and acetonitrile, respectively. Elution was performed isocratically for 2 min at 5% eluent B, from 2-13 min with linear gradient to 95% B, from 13-15 min isocratically at 95% B, and from 15.01-18 min at 5% B. The flow rate was set to 400 µL/min and the column temperature was maintained at 40 °C. Mass spectrometric analysis was performed by MS1 full scan from 65-1500 Dalton and 100 ms accumulation time (ZenoToF 7600, Sciex GmbH, Darmstadt, Germany) operating in negative mode and controlled by Sciex OS software (Sciex). The declustering potential was set to 80 (−80 V) and a spread of 50 V. MS/MS-CID fragmentation was triggered by data dependent acquisition in 20 ms pockets and up to 40 candidate spectra were recorded between 65-1500 Dalton for ions were the threshold exceeded 150 cps. For MS2 the declustering potential was set to 80 (−80 V) and a spread of 50 V, while the collision energy was set to 35 (−35 V) and a spread of 25 V. The source operation parameters were as the following: ion spray voltage, −4500 V / +5500 V; nebulizing gas, 60 psi; source temperature, 600 °C; drying gas, 70 psi; curtain gas, 35 psi CAD gas 7 psi. Instrument tuning and internal mass calibration were performed every 10 samples with the calibrant delivery system applying X500 ESI negative or positive tuning solution, respectively (Sciex GmbH, Darmstadt, Germany). The different peak areas were determined with Multiquant 3 software (Sciex GmbH, Darmstadt, Germany). Acylsugar abundances were estimated by dividing the peak area of each metabolite by the peak area of the internal standard (IS) and the leaf dry weight. 10 μM phlorizin was used as IS.

Volatile terpenoids were collected by surface extraction using leaf discs of 1 cm diameter, which were vortexed for 30 seconds in *n*-hexane. Mono- and sesquiterpenes were detected using a Trace GC Ultra gas chromatograph coupled with an ATAS Optic 3 injector and an ISQ mass spectrometer (Thermo Scientific) with electron impact ionization. The chromatographic separation was performed on a ZB-5ms capillary column (30 m × 0.32 mm, Phenomenex). The flow rate of helium was 1 ml min^−1^, and the injection temperature rose from 60 to 250 °C at 10°C sec^−1^ during 30 sec. The GC oven temperature ramp was 50 °C for 1 min, 50–150 °C at 7°C min^−1^ and 150–300 °C at 25 °C min^−1^ for 2 min. Mass spectrometry was performed at 70 eV in full scan mode with *m*/*z* from 50 to 450. Data analysis was done with the Xcalibur software (Thermo Scientific). Volatiles abundances were estimated by dividing the peak area of each metabolite to the peak area of the internal standard (IS) and the leaf dry weight. 20 μM Menthol was used as IS.

### Phytohormone quantification

OPDA, JA and JA-Ile phytohormone quantification was done as described previously for IAA (Ai et al., 2023). Extraction from 100 mg fresh weight of leaf tissue was performed by 3 consecutive cycles of Fastprep/centrifugation using 80% methanol pH 2.4 containing [^2^H_5_] OPDA, [^2^H_6_]JA, and [^2^H_2_] JA-Ile as internal standards. Phytohormone content was calculated as the ratio of analyte and internal standard peak areas, divided by the fresh weight.

The large volume separation that was used is also described in Ai et al. (2023). The chromatographic separation was done on a Nucleoshell RP Biphenyl column (2.1 x 100 mm, particle size 2.1 µm, Macherey & Nagel, GmbH, Düren, Germany) using these gradients: 0–2 min: 95% A and 5% B, 13 min: 5% A and 95% B, 13–15 min: 5% A and 95% B, and 15–18 min: 95% A and 5% B. The column was maintained at 40 °C, while the auto-sampler had a temperature of 4 °C. Solvent A was 0.3 mM ammonium formate, acidified to pH 3.0 by formic acid, and solvent B acetonitrile. Before the UPLC, 600 µl sample extract was injected into a divinylbenzene stationary-phase micro-SPE cartridge (SparkHolland B.V., Emmen, The Netherlands) at 200 µl/min and the phytohormones were trapped by adding 3,800 µl/min water and transferred from the SPE cartridge to UPLC by 120 µl 20 % acetonitrile under continuous dilution with water. The whole process was done on a prototype system consisting of CTC Combi-PAL autosampler equipped with a 1 ml injection loop, an ACE 96-well plate SPE unit, a high-pressure dispenser, a SPH1299 UPLC gradient pump, an EPH30 UPLC dilution pump, and a Mistral column oven (all from AxelSemrau GmbH, Sprockhövel, Germany).

Phytohormone level estimation after short HS was performed according to (Balcke et al., 2012) using 50 mg of homogenized plant leaf material. The extraction was done in 250 μL of pure methanol. After full speed centrifugation, the supernatant was diluted using 1750 μL of water and the extracts submitted to solid phase extraction (SPE) on HR-XC column (Macherey Nagel Filterplates). Samples were eluted in 900 μL acetonitrile and concentrated in speedvac for 1 h. Phytohormone separation was performed similar to the separation of semi polar metabolites described above. Only here, the following gradients were used: isocratically 2 min at 5% B, from 2-19 min linear gradient to 95% B, from 19-21 min isocratically at 95% B and from 21.01 min-24 min at 95% B.

Mass spectrometric analysis of both hormone separations was performed by scheduled multiple reaction monitoring (MRM) on a QTRAP6500 (Sciex GmbH, Darmstadt, Germany) operating in negative ion mode and controlled by Analyst 1.7.1 software (Sciex GmbH, Darmstadt, Germany). The source operation parameters were as the following: ion spray voltage, −4500 V / +5500 V; nebulizing gas, 60 psi; source temperature, 450 °C; drying gas, 70 psi; curtain gas, 35 psi.

Relative abundances of the phytohormones from the short HS experiments were calculated by dividing the peak area to the fresh weigh.

### Arabidopsis growth conditions and heat treatment

Seeds of *Arabidopsis thaliana* Col-0 were germinated on soil. 10-day-old seedlings were transferred to individual pots filled with steam-sterilized soil and maintained in growth chambers (Percival Scientific, Perry, IA, USA) under short day conditions and controlled temperature (18 °C/18 °C) and humidity (65–70%) for 10 days. Control (non-stressed plants) were kept under the same conditions, whereas heat stress was applied by growing plants at 37 °C at day and 23 °C at night. After 16 days, leaves number 8 and 13 were harvested and photos taken using a Zoom microscope (AZ100, Nikon). Trichome number was determined per leaf using ImageJ.

## Acknowledgments und funding

This work was funded by grant # TI 800/10-1 from the *Deutsche Forschungsgemeinschaft* and grant SAW-2020-IGZ-1.1.1.1-Ketchup from the Leibniz Association to AT. We thank Hagen Stellmach for assistance in the hormone measurements and the gardeners of the Leibniz Institute of Plant Biochemistry for plant care.

## Author contributions

This project is based on an initial observation by NB that heat stress induces type-IV trichomes in tomato. AT designed and supervised the project, and wrote part of and edited the manuscript. RS and AB performed the experiments, prepared the figures and wrote part of the manuscript. GUB supervised LC-MS measurements. MP-R and BH performed the Arabidopsis heat stress experiments. The environmental scanning electron microscopy images were produced by FS. All authors read and approved the manuscript.

---

**Supplemental Figure 1.**
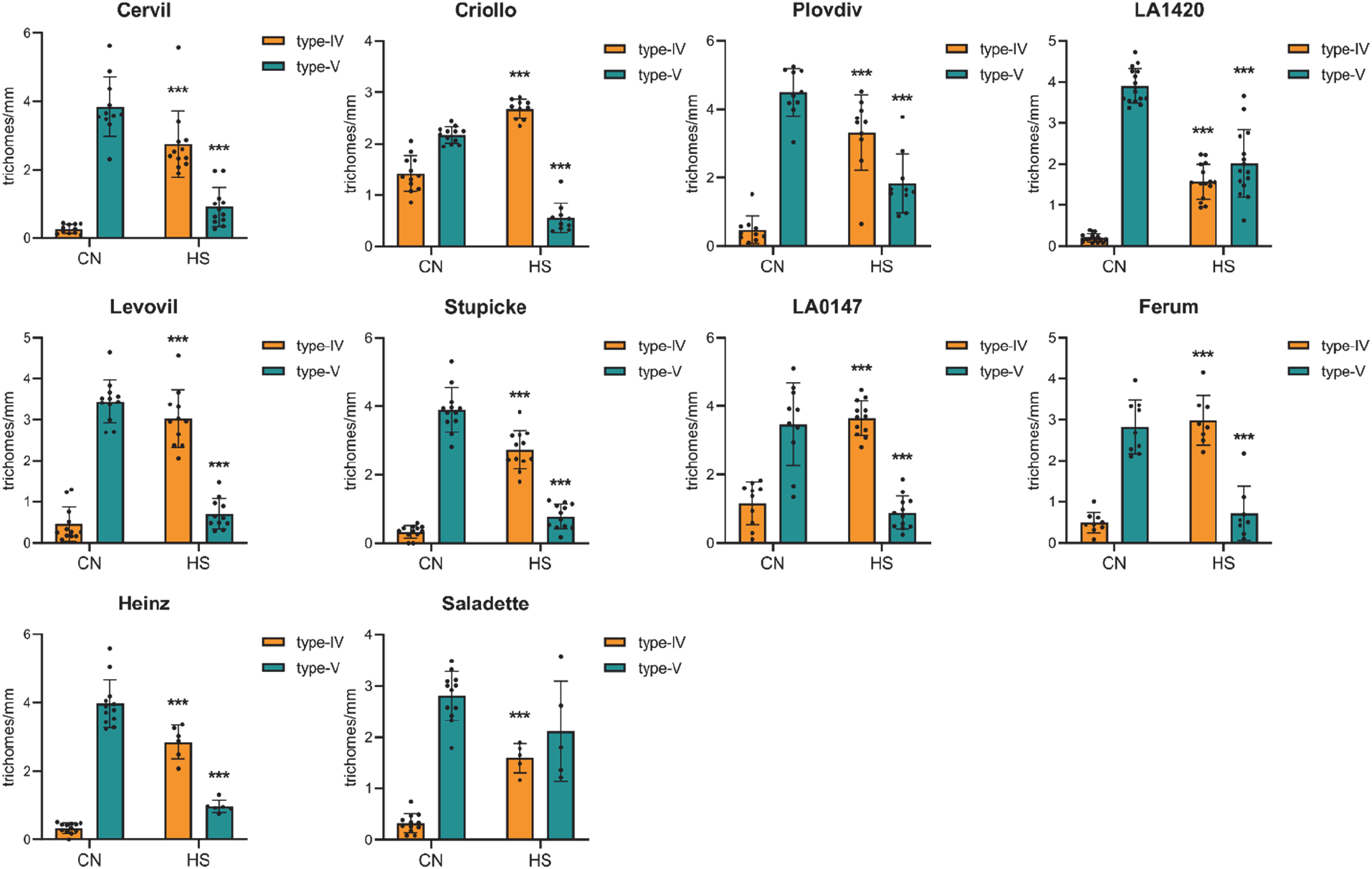
Trichome counting from the edge of leaves of different tomato varieties after three weeks of growth under CN and HS conditions. Each dot indicates a biological replicate. Error bars indicate standard deviation. Pairwise comparisons were done using Student’s *t*-test \**P* < 0.05, \*\**P* < 0.01, \*\*\**P* < 0.001. *P* values indicated in relation to corresponding CN.

**Supplemental Figure 2.**
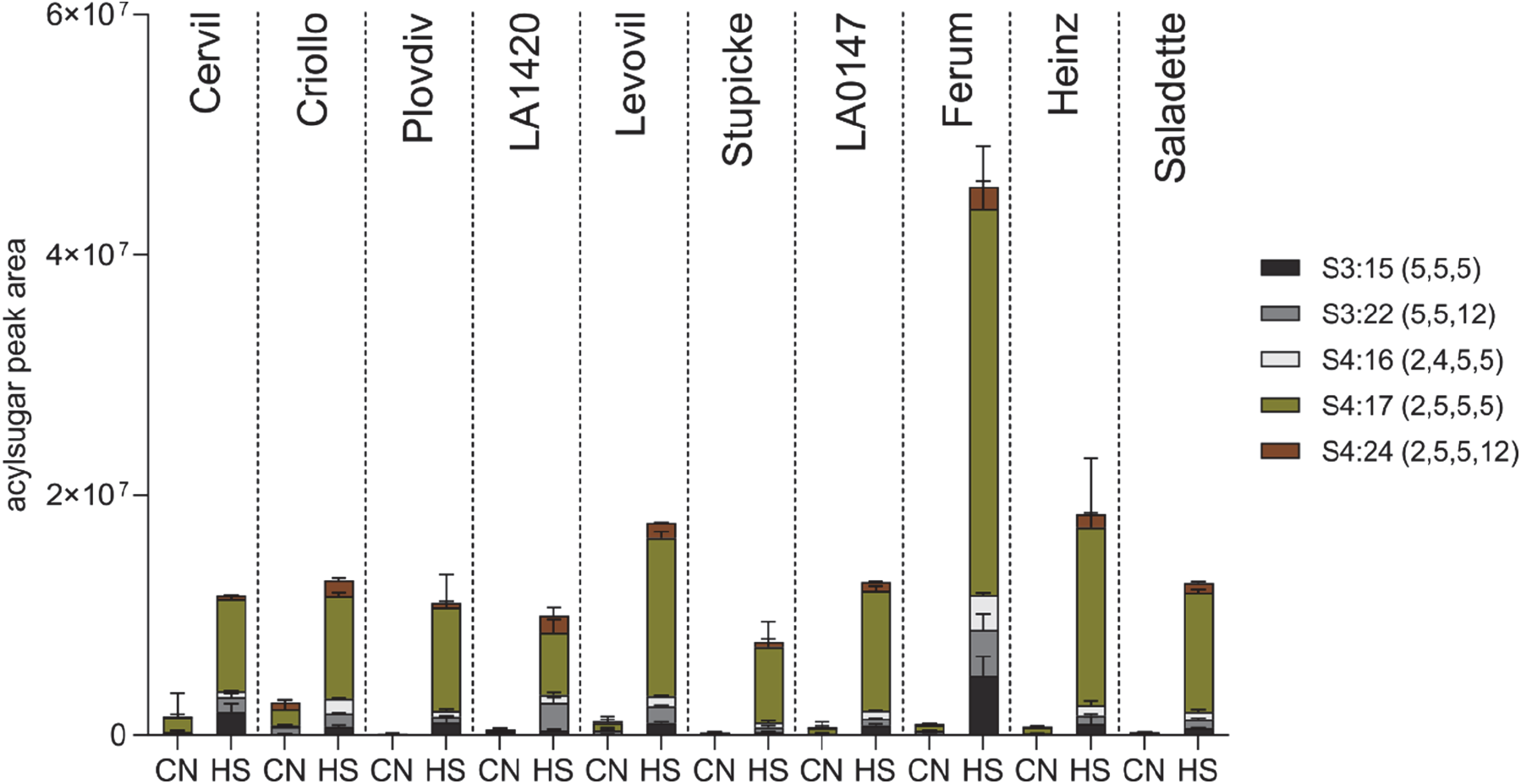
Liquid chromatography-mass spectrometry (LC-MS) quantification of acylsugars of plants after 3 weeks of growth under CN and HS conditions (*n* = 3 biological replicates). Error bars indicate standard deviation.

**Supplemental Figure 3.**
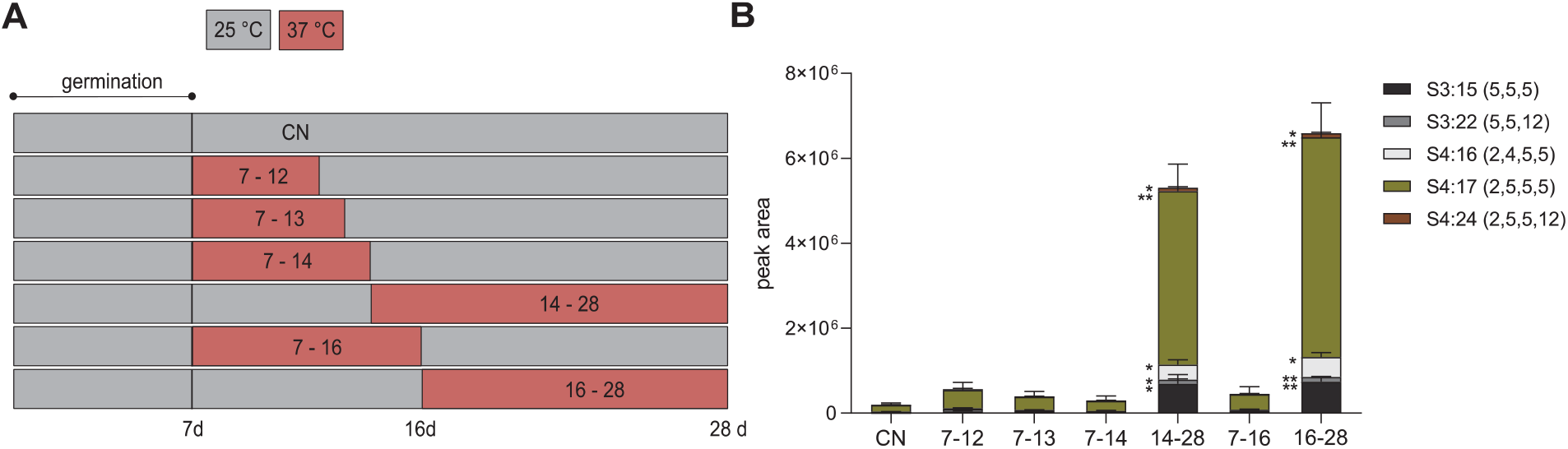
Time regime (A) and quantification of AS (B) from plants stressed with different heat periods (n = 3 biological replicates). Error bars indicate standard deviation. Pairwise comparisons between HS and CN were done using Student’s *t*-test \**P* < 0.05, \*\**P* < 0.01, \*\*\**P* < 0.001.

**Supplemental Figure 4.**
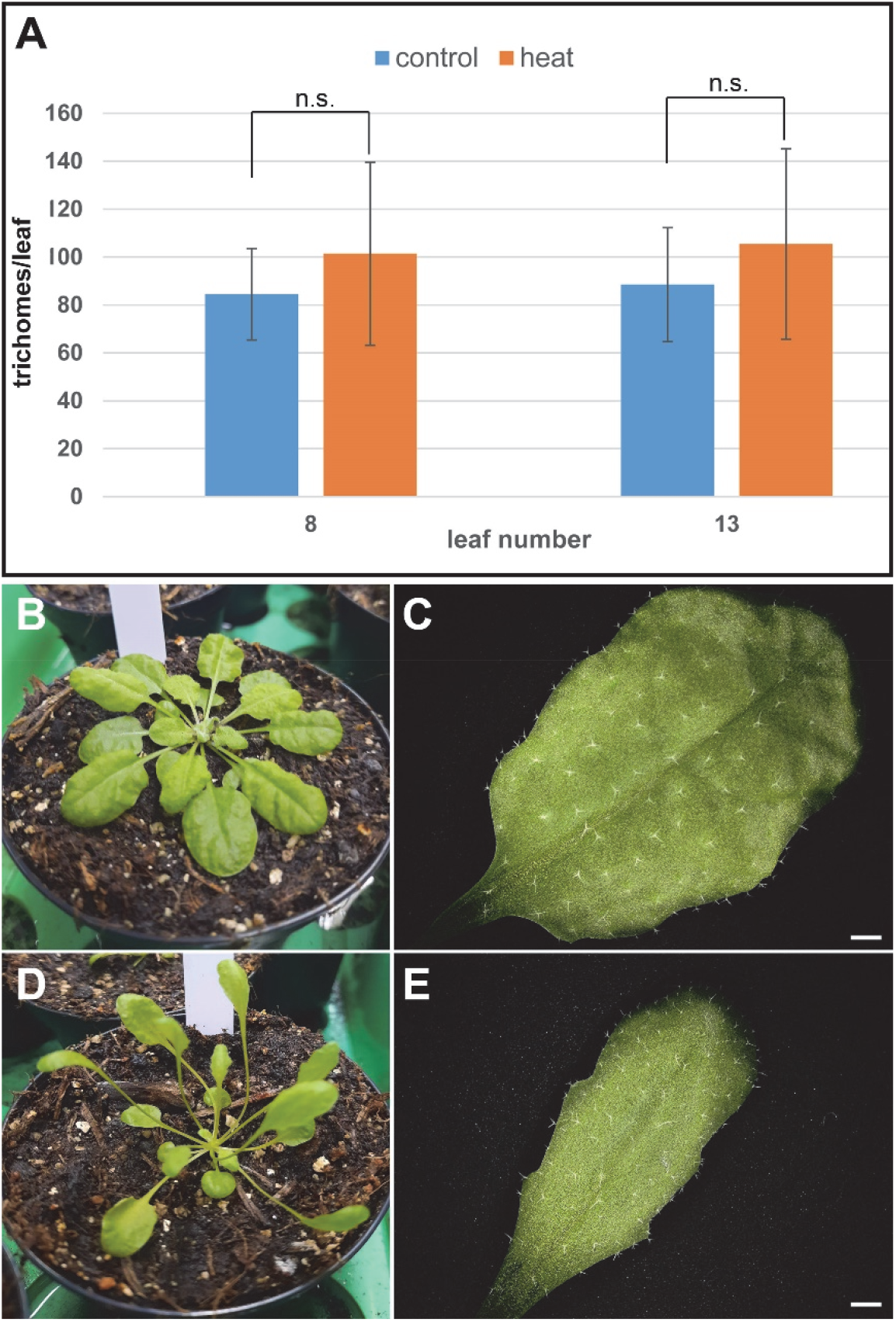
Trichome numbers and plant phenotype of *Arabidopsis thaliana* after heat stress. A. Trichome numbers of leaf 8 and leaf 13 of plants grown under normal (control) and heat conditions (day temperature 37 °C) for 16 days. Each value is represented by the mean of ten biological replicates ± SD. Treatments and controls were pairwise compared by the Student’s t-test and did not reveal significant differences (n.s.). B, C. Plant and leaf phenotype of plants grown under control conditions. D, E. Plant and leaf phenotype of plants grown under heat conditions. Note the typical phenotype of heat-stressed plants visible by elongated petioles, leaf hyponasty and a reduction in leaf blade size. Bars in C and E represent 1 mm.

**Supplemental Table 1.**
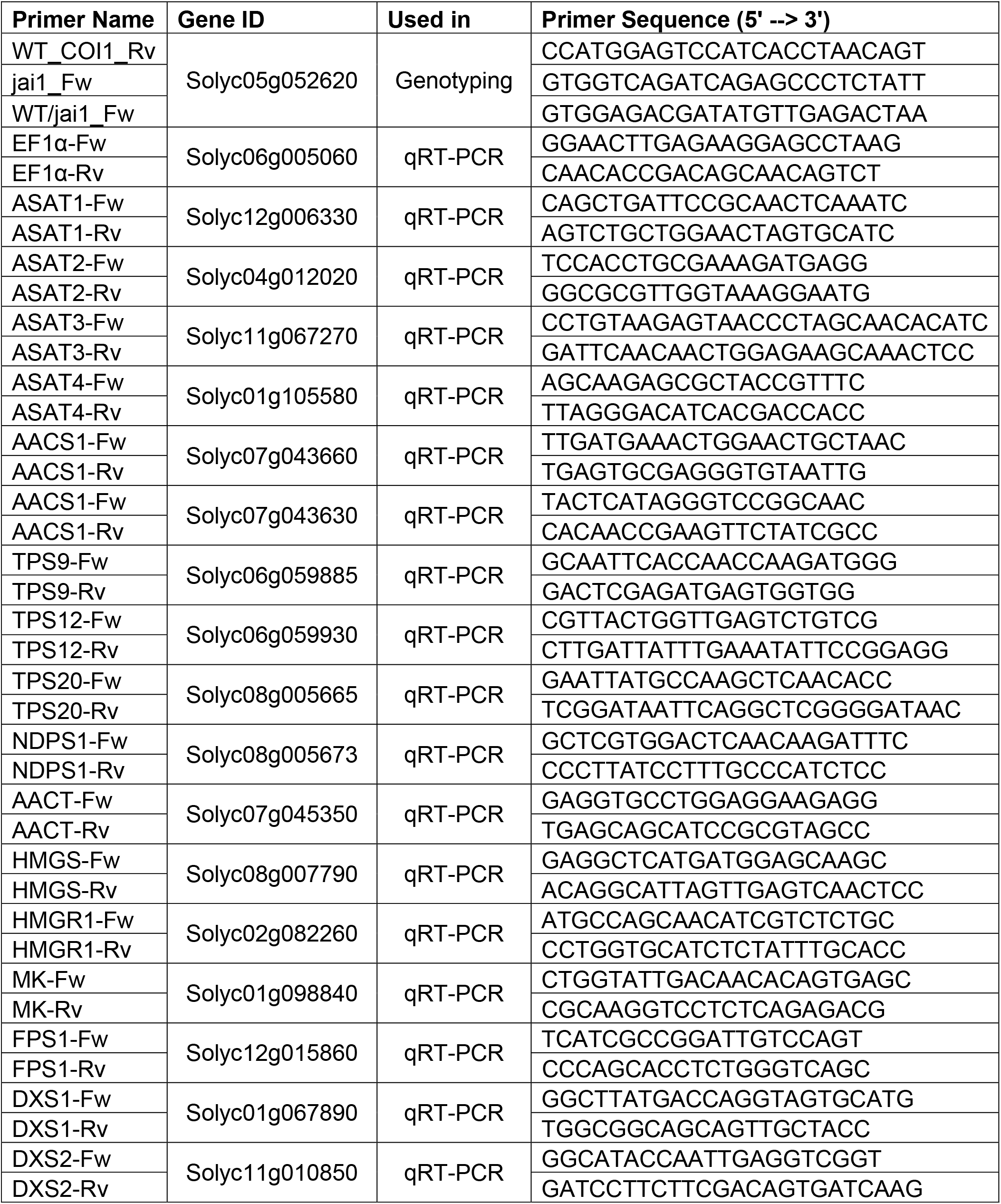
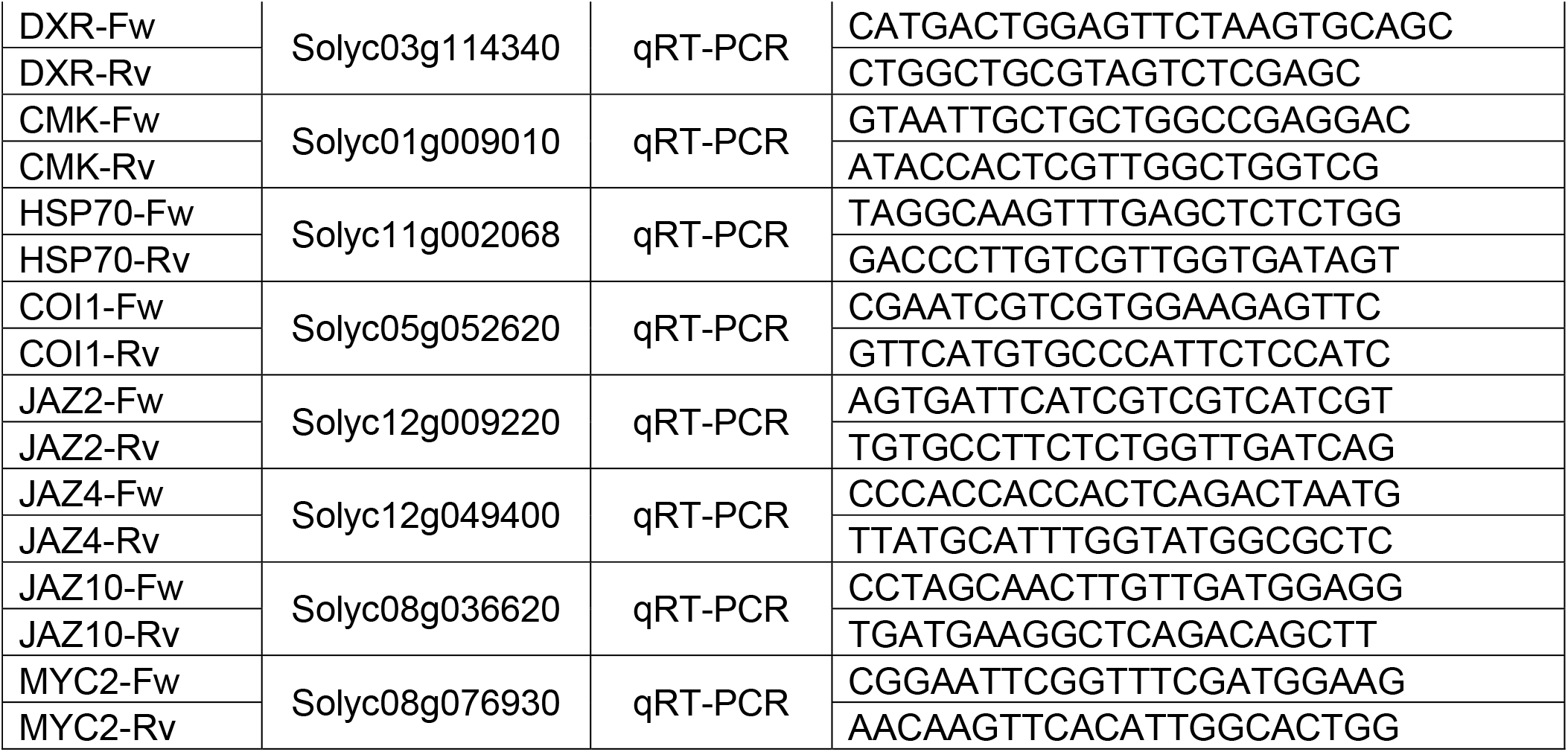
Primers used in the present study.

